# Evolution of reproductive isolation in a long-term evolution experiment with *Drosophila melanogaster*: 30 years of divergent life history selection

**DOI:** 10.1101/2023.02.01.526683

**Authors:** Chloe E Robinson, Harshavardhan Thyagarajan, Adam K Chippindale

## Abstract

We ask if three decades and over 1 500 generations of divergent life history selection on age at reproduction has resulted in the evolution of reproductive isolation (RI) between laboratory populations of *Drosophila melanogaster*. We tested for premating, postmating-prezygotic and postzygotic reproductive isolation between 3 replicate population pairs. Large evolved differences in body size between selection treatments suggested the potential for prezygotic barriers driven by sexual selection or physical incompatibilities between the sexes. Although a simple prediction would be preference for larger size, creating directional isolation, our results from individual mate choice trials indicate that populations from both selection treatments show a marked bias towards homotypic mate choice; indicative of prezygotic RI driven by sexual selection or sexual conflict. Hybridization between the focal populations resulted in the production of viable adult flies with intermediate size and developmental traits. We observed a suggestive but statistically non-significant trend of fitness decline in the F2 generation of hybrids, but no significant evidence suggesting the evolution of postmating-prezygotic or postzygotic RI. Our findings are in accord with extant literature that posits that premating RI evolves before postmating forms of RI.

## INTRODUCTION

Understanding the mechanisms underlying the evolution of reproductive isolation and the timing of their origination is central to the study of speciation. In this work, we focus on the role of divergent life-history selection in ecological speciation. Ecological speciation is the process through which reproductive isolation evolves as a consequence of local adaptation with limited gene flow. Divergent selection pressures can contribute to this process by acting directly, or through pleiotropy or tight linkage with genes that are directly responsible for reproductive isolation (Rice and Hostert 1993; Rundle and Nosil 2005). Divergent evolved responses between ecotypes may result in (1) changes in morphology, mating behaviours or physiological characters that hinder mating attempts mechanically or via mate choice (premating reproductive isolation); (2) changes in gametes/reproductive machinery that inhibit fertilization (postmating-prezygotic reproductive isolation); and/or (3) genetic incompatibilities that reduce survival or reproductive success of hybrid individuals (postzygotic reproductive isolation).

In addition to divergent natural selection in allopatric populations, divergent intersexual coevolution via sexual selection has long been recognized to have a potentially significant role in the evolution of reproductive isolation (Lande 1981; Maan and Seehausen 2011; Panhuis *et al*. 2001; Ritchie 2007; Singh and Singh 2014). Likewise, interlocus sexual conflict – where traits that increase the fitness of one sex directly cost the other – can result in rapid coevolution (an “evolutionary chase” or “arms race”) between male and female reproductive traits (Rice & Holland 1997; Parker & Partridge 1998; Rice 1998) and potentially accelerate the process of speciation (Ritchie 2007; Gavrilets 2014; Syed *et al*. 2017); the evolution of female resistance to reduce the direct costs of mating is predicted to contribute to assortative mate choice (Gavrilets *et al*. 2001).

While natural systems such as hybrid zones have advanced our understanding of the evolution of RI immensely, researchers are frequently limited to indirect inferences, such as using estimates of gene flow and clines to infer reproductive isolation. Experimental evolution can be a valuable tool for more direct investigation, allowing us to probe potential drivers of speciation between populations with well understood evolutionary histories. Experimental studies in speciation have attempted to test if (1) non-ecological drivers like drift, (2) negative selection against hybridization between ecotypes, (3) divergent selection on populations or (4) sexual conflict produced reproductive isolation (reviewed in Rice and Hostert 1993; Coyne and Orr 2004; Fry 2009; White *et al*. 2020). An obvious limitation to experimental evolutionary studies of speciation is time: research projects rarely exceed granting cycles and few organisms lend themselves to experimental evolution. It is little surprise then that *Drosophila* is far and away the genus most investigated in the laboratory.

Early divergent selection studies produced seemingly equivocal results with respect to creating reproductive isolation. A closer examination suggested that divergent selection was highly effective, given a pre-existent tendency to mate assortatively on the basis of the trait selected upon (Rice and Hostert 1993; Fry 2009). Divergent selection upon behavioural traits resulted in the correlated evolution of reproductive isolation (del Solar 1966; Soans *et al*. 1974; Hurd and Eisenberg 1975; Lofdahl *et al*. 1992) while divergent selection on other traits like bristle number (Barker and Cummins 1969; Santibañez and Waddington 1958) produced negative results. More recently, experimental evolution studies have focused on selection regimes that do not directly involve characters that are known to influence intra-population assortment. For example, a number of studies have investigated the importance of local adaptation to distinct nutrition sources for mating assortment. Divergent selection regimes with different nutritional environments result in premating isolation through (1) the evolution of signalling traits and mating preferences in *Drosophila serrata* (Rundle et al. 2005), (2) the evolution of symbiotic microbiota in inbred strains of *D. melanogaster* (Najarro *et al*. 2015; Sharon *et al*. 2010), (3) competitive ability differences between controls and selected individuals in *D. melanogaster* (Belkina *et al*. 2018), (4) positive assortative mating preferences (albeit unstable) in *D. melanogaster* (Nash *et al*. 2019); and (5) postzygotic reproductive isolation through genetic incompatibilities in *Saccharomyces cerevisiae* (Dettman *et al*. 2007) and *Neurospora* (Dettman *et al*. 2008).

Life history traits are less frequently studied in the context of reproductive isolation, but have also received some attention. For example, divergent selection on pre-adult development time has been shown to produce pre-mating isolation through (1) disturbances in circadian rhythm resulting in changes in phases of mating in the melon fly, *Bactrocera cucurbitae* (Miyatake and Shimizu 1999), and (2) evolved body size differences, and correlated levels of sexual conflict, in *Drosophila melanogaster* (Ghosh and Joshi 2012). Likewise, divergent selection on body size results in premating isolation, through mechanical incompatibilities during attempted copulations between *Columbicola* feather mites (Villa *et al*. 2019).

Age-at-reproduction has been extensively manipulated using experimental evolution in *Drosophila* to investigate correlated changes in longevity, development time, body size and a slew of associated traits (Prasad and Joshi 2003). In this study, we use an exceptionally long term evolution experiment (LTEE) with population pairs separated by over 1,500 generations and three decades under divergent selection in allopatry. The two *D. melanogaster* life-history selection regimes used in this study are selected to reproduce late in life (4 weeks from egg; CO) and extremely early in life (9 days from egg; Accelerated CO / ACO). The ACO populations have evolved small body size, rapid development time, and reduced lifespan (Chippindale *et al*. 1997a; Chippindale *et al*. 2004a). On the other hand, the CO flies are big, develop slowly and likely have an increased ability to resist copulations as females and to inflict harm as males (Chippindale *et al*. 2004b; Chippindale *et al*. 1997b; Mital *et al*. 2021, Verma *et al*. 2022).

In the speciation literature, body size is identified as a crucial trait that can drive the evolution of reproductive isolation (Servedio et al. 2011). In *D. melanogaster*, body size is strongly correlated with juvenile development time (Chippindale *et al*. 1997a; Nunney 1996; Prasad and Joshi 2003; Prasad *et al*. 2000; Zwaan *et al*. 1995) and life-history selection on development has been shown to result in the evolution of body size (Chippindale *et al*. 1997a; Prasad *et al*. 2000). An important sexually selected trait for both sexes, large females tend to be more fecund (Roff 2002; Stearns 1992) and attract higher courtship effort from males (Byrne and Rice 2006); while large males have increased competitive abilities (Markow and Ricker 1992; Partridge and Farquhar 1983), and secure more frequent copulations as a consequence of female preference (Markow 1986; Partridge *et al*. 1987, but see Prasad et al. 2007). Given the evolved size differences between the ACOs and COs, a simple prediction is that individuals of both population-types display a mating preference for individuals from the larger CO populations, producing conditions of partial, directional, reproductive isolation driven by sexual selection on the basis of body size, along the lines observed by Ghosh and Joshi (2012).

In *D. melanogaster*, divergent reproductive strategies between the sexes have long been recognized (Bateman 1948) potentially resulting in interlocus sexual conflict. Frequent matings increase male fitness but can decrease female fitness as a result of male harassment and the harmful effects of genital wounding and seminal fluid (Chapman et al. 1995; Bonduriansky *et al*. 2008; Travers *et al*. 2015). The harm experienced by females as a result of mating, measured by lifespan and egg production rate, is correlated with male body size (Pitnick and García–González 2002; Friberg and Arnqvist 2003). In the context of our two selection treatments, we predict that male harm and female resistance traits will be stronger in the CO populations compared to the ACO populations. In experimental hybridization between CO and ACO lines, this difference could drive premating isolating barriers to evolve in one direction, with CO females resistant to mating with the smaller ACO males, and ACO females vulnerable to unwanted mating with CO males. Indeed, working with a similar system, Ghosh and Joshi (2012) found such one-directional isolation, driven by large females resisting mating attempts by small males, and a preference for (or lack of resistance to) large males in females of both treatments. On the other hand, especially with the large number of generations in allopatry, mating signals and preferences may have diverged between populations in our selection experiment creating the possibility for positive assortment in mate choice experiments.

We investigated mate choice through individual female, male, and group mating choice assays in the ACO/CO complex of selected lines. We further looked for potential gametic incompatibilities (postcopulatory prezygotic isolation) by comparing egg hatchability in hybrid matings to the parental populations. Finally we looked for postzygotic effects (hybrid breakdown or vigour) in developmental traits, body size, and adult fitness.

## METHODS

### Experimental populations

Our experiments employed the accelerated development (ACO) (Chippindale *et al*. 1997a) and control-old (CO) (Rose et al. 1992) populations. All populations are maintained under standard laboratory conditions: moderate density (larval density of 80 - 120 larvae / vial), 25ºC, 12:12 light:dark cycle on banana/agar/killed-yeast medium, and with a census population size of approximately 2,000 individuals per generation.

The five CO populations (CO_1_-CO_5_) were derived from like-numbered O populations in 1989 (Rose *et al*. 1992). Each CO population is maintained on a 28-day discrete life cycle, with flies placed into cages at a density of ∼1000 flies/cage. On day 26, yeast paste is added to the medium to stimulate egg-laying, and on day 28/day 0, eggs are collected at 80-120 eggs per vial. In 1991, one A population (ACO_1_-ACO_5_) was derived from each of the five CO populations and initially subjected to direct selection for accelerated development. After generation 175, these accelerated populations were maintained on a discrete life cycle of 9 days.

Experimental populations were derived from three of the five pairs of ACO and CO populations (ACO_1_, ACO_3_, ACO_5_ and CO_1_, CO_3_, CO_5_). To remove environmental effects and synchronise experiments, all populations were maintained over a control 14 day life cycle for two generations prior to any assay. Prior to experiment, life cycles were staggered in light of evolved development time differences (ACO << CO) with a 48h difference in egg collection time to synchronise eclosion time of adults. Each assay was repeated for three replicate population pairs: CO_1_-ACO_1_, CO_3_-ACO_3_, and CO_5_-ACO_5_.

#### Female mate choice assay

To test for the existence of premating reproductive isolation between populations as mediated by female preference, we evaluated the mate choice of ACO and CO females. Flies were sorted by sex and collected as virgins within 6 hours of eclosion under light CO_2_ anaesthesia. A food dye technique was used to differentiate between males from each regime (developed from Verspoor *et al*. 2015). Dye techniques were calibrated based on tests of mortality, and the volume of dye used did not result in detectable harmful side-effects, measured through changes in lifespan. Vials were dyed three days prior to the assay using 6 drops of red or blue food dye. One day prior to the experiment, male flies were added to coloured vials at a density of 10 flies/vial and left to ingest the dye for 24 hours. Females were acclimated to individual vials for at least 24 hours prior to the assay. At the time of the assay one male of each population identity and colour was mouth aspirated into an empty vial, then flipped into a vial containing an acclimatised female. Mating latency (time to amplexus formation), colour of the mated male and mating duration were recorded. Under the conditions we used, copulations lasted on average between 15-20 minutes. A viable mating was considered one that lasted for more than two minutes, and vials in which mating did not occur within 60 minutes were discarded. This represents a commonly employed design to test mate choice (Dukas 2005, Ghosh & Joshi 2012, Arbuthnott *et al*. 2017). Vials were randomised so that observers were unaware of female identity. Trials were colour balanced so that in half of the trials the ACO male was blue and the CO male red, and in the other half the alternate dye pattern was applied. Flies were 2-3 days old at the time of the assay.

This assay was repeated over two days, with female choice tested in both a morning (9am-12pm) and afternoon (1pm-4pm) trial each day. In each of the 4 trials, we tested the choice of 50 females from both selection regimes for each of the 3 replicate populations. The ratio of homotypic to heterotypic matings was calculated across the 50 replicate vials for each combination of replicate population and selection regime.

#### Male mate choice assay

It is difficult to distinguish the influence of male competition and courtship effort from female preference and resistance. By testing individual male mate choice in addition to female choice and contrasting the results, we hoped to uncover the source of any observed non-random mating pattern. From the time of virgin collection, males were maintained at 10 males/vial. Two females, one from each selection regime, were acclimatised to the same vial 24 hours prior to the assay. Female flies from ACO and CO selection treatments could be identified reliably based upon size alone. On the day of the experiment, one male was “pootered” (mouth aspirated) into a vial containing two females. As with the female choice assay, observers recorded mating latency, the size of the mated female (large or small) and mating duration and all other procedures were the same.

#### Group mate choice assay

To test for the existence of premating reproductive isolation between ACO and CO populations when mating occurred in a group environment, we evaluated female mate choice in vials containing 10 females and 12 males from each regime. Flies were handled and marked using the procedures outlined above for individual assays. On the day of the assay, male vials of paired ACO and CO populations were combined and then flipped into a female vial. Vials were observed until at least 8 (out of a possible 10) simultaneously copulating pairs were observed, and were then frozen for 60 minutes without disruption to the copulating pairs. The frozen pairs were observed and the colour of each mated male was recorded. This methodology was repeated over two trials. During each trial, 10 vials were tested for each regime within each replicate population.

#### Hatchability

In order to determine the existence of postmating prezygotic isolation we compared the hatch rate of eggs produced from F1 and F2 crosses CO_*i*_♀ × ACO_*i*_♂and ACO_*i*_♀ × CO_*i*_♂ as well as the parental crosses CO_*i*_♀ × CO_*i*_♂, and ACO_*i*_♀ × ACO_*i*_♂ (*i* = 1,3,5). Crosses were performed in vials with 12 virgin females and 10 virgin males. A period of at least 4 hours was allowed for mating and egg laying. From each cross, male and female progeny were collected as virgins and maintained in sex specific vials. Hybrid and parental crosses were staggered in an attempt to synchronise time of eclosion.

For tests of hatchability, 100 flies of each sex were deposited in a collector bottle with a plate of yeasted food. After an egg laying period of 3 hours the plate was removed. Eggs were transferred into vials using fine brushes and egg collection solution (Ashburner 2005). Each vial contained 90 eggs arranged in a grid pattern to facilitate easy counting. 9 such vials were created for each cross identity, for each of the three replicate populations. 24 hours after egg collection, the proportion of hatched eggs was determined. Vials continued to be maintained under standard conditions to be used in tests of viability, development time, and size.

#### Larvae to adult viability

To uncover any survival differences, we compared the larvae to adult viability of hybrids compared to flies from parental crosses. Twelve days after egg collection, the number of eclosed adults in each vial was recorded. The difference between the number of viable eggs and eclosed adults was used as a measure of larvae to adult viability. 9 vials were tested for each cross identity in each replicate population.

#### Development time

Development time of flies from hybrid and parental crosses was also analysed as an indicator of hybrid viability. Each vial was observed for eclosion over a period of 4 days, beginning 24 hours prior to the time of expected peak eclosion for each line. For the first 2 days, vials were observed every 6 hours. For the following 2 days, observations were made every 12 hours. During each observation, the number of eclosed flies was recorded. An average eclosion time was then calculated for the vial. 9 vials were tested for each cross identity across each replicate population.

#### Body Size

During peak eclosion, 20 female flies were collected from each cross identity. These flies were given 1 hour to physically mature and then were frozen. Flies were divided into 4 groups of 5 females and placed into tinfoil dishes to be dried in an oven at 70 degrees Celsius for 24 hours. Each dish was measured in a microbalance four times: twice with the flies and twice without. These totals were averaged and divided by 5 to provide an estimate of the dry weight of an individual fly.

#### Fertility - Competitive reproductive fitness

An adult fitness assay was completed to test for parental line and hybrid fertility differences. In this assay, red-eyed focal flies competed with control flies marked with a recessive brown-eye marker (*bw*^1^) derived from the baseline. The IVb population was selected as these flies are a common ancestor to both the focal populations, also having an intermediate life-history selection protocol. The recessive nature of *bw* allowed the proportion of red eyed progeny to be used as a metric of paternity share.

Focal F1 and F2 flies of the hybrid crosses CO_*i*_♀ × ACO_*i*_♂, ACO_*i*_♀ × CO_*i*_♂ as well as the parental crosses CO_*i*_♀ × CO_*i*_♂, and ACO_*i*_♀ × ACO_*i*_♂ (*i* = 1,3,5) were tested. Crosses were performed in vials with 12 virgin females and 10 virgin males. A period of at least 4 hours was allowed for mating and egg laying. From each cross, male and female progeny (focal flies) were collected as virgins and maintained in sex specific vials at a density of 10 flies/vial.

Focal flies were anaesthetized under CO_2_ and combined with four IVb flies of the same sex (competitors) and four IVb flies of the opposite sex (potential mates). Vials were supplemented with a sparse amount of yeast. Female focal and IVb flies were acclimatised to vials 24 hours prior to the assay. After 48 hours, flies were flipped into new vials where oviposition occurred. Flies were 5-7 days old at the time of the assay. After 18 hours flies were discarded. At the time of eclosion, the number of red eyed and brown eyed offspring produced from the competition assay were counted. Thirty vials for each sex and cross identity were tested for each of the 3 replicate populations.

#### Statistical Analysis

Statistical analysis was conducted in RStudio version 1.3.1073 (RStudio Team 2020) and R version 1.3.1073 (R Core Team 2020). Data was visually assessed for residual normality and heterogeneity. Female, male and group mate choice data was analysed with a repeated G-test for goodness of fit (Ghosh & Joshi 2012, McDonald 2014). In each case, the ratio of homotypic to heterotypic matings was tested for deviation from a 1:1 null expectation separately for each replicate population and overall for each selection regime. To determine the effect of cross identity on fly development time and body size, linear mixed effects models (LMMs) were fit with cross identity as a fixed effect and replicate population as a random effect. To determine the effect of cross identity on hatchability, viability and the fertility of flies, binomial generalised linear mixed effects models (GLMMs) were fit with cross identity included as fixed effect and parental replicate included as a random effect. Fertility data was analysed separately for males and females. Models were fit using the ‘*lme4*’ package (Bates et al. 2015). Fixed effects and random effects were analysed using the ‘*anova()*’ and ‘*ranova()*’, and ‘*Anova()*’ and by backwards model selection for LMMs and GLMMs, respectively (Kuznetsova et al. 2017, Fox & Weisberg 2019). Post-hoc comparisons between cross identities were performed using the ‘*emmeans*’ package (Lenth 2022).

## RESULTS

### Female Mate Choice Assay

Overall, ACO females showed a significantly higher proportion of homotypic (0.57 +/- 0.08, mean +/- sd) compared to heterotypic matings (0.42 +/- 0.08, *P* = 0.01, Figure 1a), with no significant heterogeneity among replicate population pairs (*P* = 0.66, Table S1). The ACO_*5*_ population showed a significant deviation from the null hypothesis, with more homotypic matings compared to heterotypic matings (*P* = 0.04; Figure S1a). Females from the ACO_*3*_ population showed a similar trend towards more homotypic matings with borderline significance (*P* = 0.05). ACO_*1*_ females showed no significant difference between mating types (*P* = 0.26). When data was pooled across trials within each replicate population, both ACO_*3*_ and ACO_*5*_ populations, but not the ACO_1_ population, showed a significant deviation from a random mating ratio (*P*_*1*_ = 0.21, *P*_*3*_ = 0.02, *P*_*5*_ = 0.02). with no significant heterogeneity between individual trials for any of the 3 replicate populations (*P*_*1*_ = 0.30, *P*_*3*_ = 0.28, *P*_*5*_ = 0.21).

**Figure 1.**
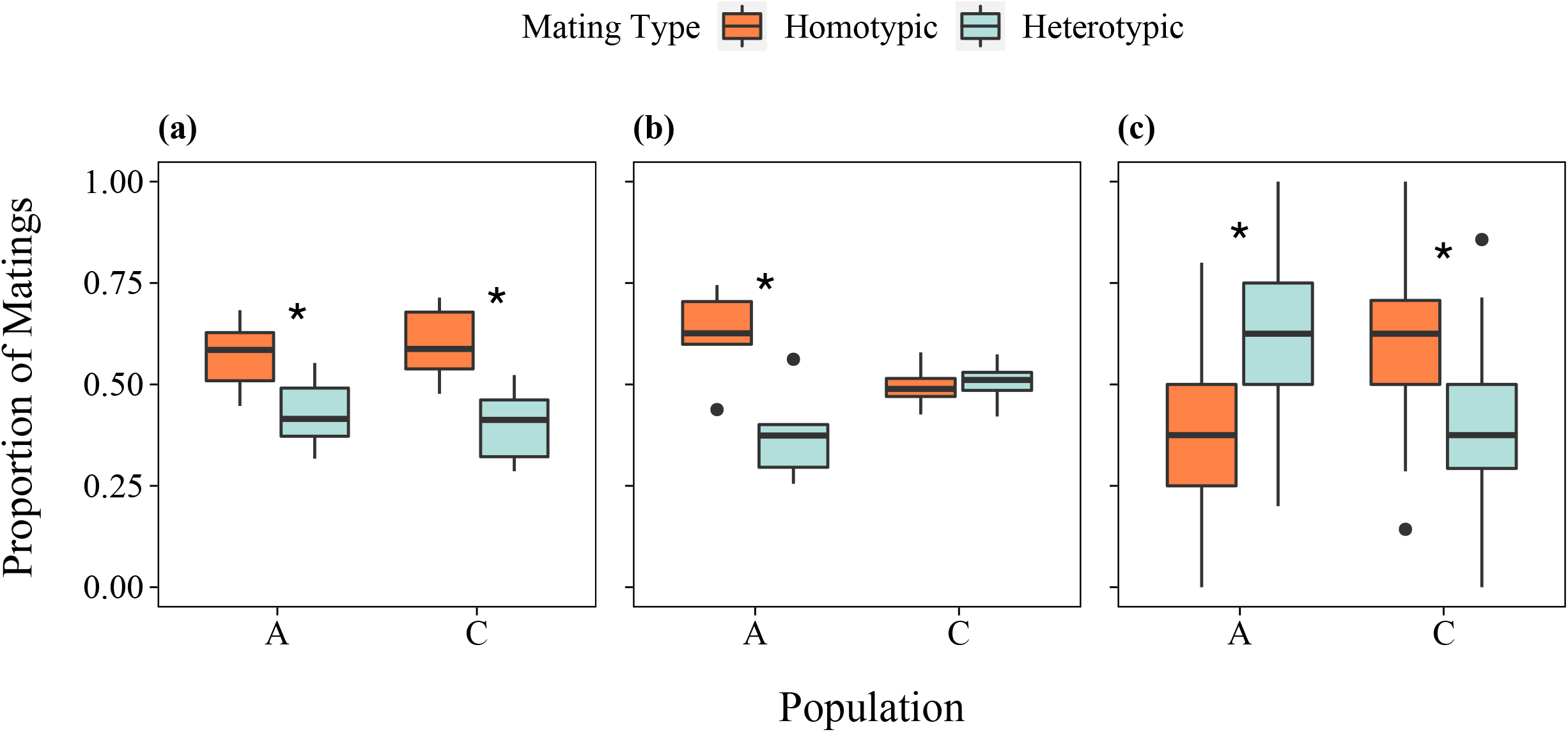
The proportion of homotypic compared to heterotypic matings recorded during the female (a), male (b) and group (c) mate choice assay. Female (a,c) and male population identity is shown on the x-axis. An asterisk represents a significant deviation from a random mating ratio as indicated by a repeated G-test for goodness of fit (p<0.05).

Overall, CO females showed a significantly higher proportion of homotypic (0.60 +/- 0.08) compared to heterotypic matings (0.40 +/- 0.08, *P* = < 0.0001, Figure 1a), with no significant heterogeneity among replicates (*P* = 0.65, Table S1). Two CO populations, CO_*1*_ and CO_*3*_, showed an overall significant deviation from the null hypothesis, with more homotypic copulations (*P*_*1*_ = < 0.01, *P*_*3*_ = 0.01, Figure S1a). CO_*5*_ females showed a trend towards more homotypic matings, but this was not significant (*P* = 0.08). When data was pooled across trials, all three CO populations showed a significant deviation from the null hypothesis of random mating (*P*_*1*_ = < 0.001, *P*_*3*_ = 0.01, *P*_*5*_ = 0.04), with no significant heterogeneity among trials for any of the replicate populations (*P*_*1*_ = 0.13, *P*_*3*_ = 0.15, *P*_*5*_ = 0.28).

### Male Mate Choice Assay

Overall, ACO males showed a significantly higher proportion of matings with ACO females (0.64 +/- 0.08) compared to CO females (0.36 +/- 0.08, *P* = < 0.00001, Figure 1b), with no significant heterogeneity between populations (*P* = 0.81, Table S2). All three A populations showed an overall significant deviation from the null hypothesis, displaying more homotypic compared to heterotypic matings (*P*_*1*_ = < 0.0001, *P*_*3*_ = 0.01, *P*_*5*_ = < 0.001). There was significant heterogeneity observed among trials within the ACO_*1*_ population (*P* = 0.01), but not the ACO_*3*_ (*P* = 0.70) or ACO_*5*_ (*P* = 0.38) populations.

Overall, CO males displayed a nearly equal proportion of matings with ACO (0.51 +/- 0.04) and CO females (0.49 +/- 0.04, P = 0.95, Figure 1b), with no heterogeneity among replicate populations (P = 0.93, Table S2). All three CO populations showed no significant deviation from the null hypothesis of random mating (*P*_*1*_ = 0.99, *P*_*3*_ = 0.65, *P*_*5*_ = 0.96). There was also no significant heterogeneity among trials within any of the replicate populations (*P*_*1*_ = 0.97, *P*_*3*_ = 0.54, *P*_*5*_ = 0.91).

### Group Mate Choice Assay

Overall, ACO females showed a significantly higher proportion of matings with CO males (0.62 +/- 0.20) compared to ACO males (0.38 +/- 0.20, *P* = <0.00001, Figure 1c), with significant heterogeneity being observed between replicate populations (*P* = 0.01, Table S3). ACO_*3*_ and ACO_*5*_ females showed an overall significant deviation from the null hypothesis, displaying more heterotypic compared to homotypic matings (*P*_*3*_ = 0.02, *P*_*5*_ = <0.0001). ACO_*1*_ females showed no deviation from the null expectation (*P* = 0.78). There was no significant heterogeneity observed among trials within any of the three replicate ACO populations (*P*_*1*_ = 0.49, *P*_*3*_ = 0.45, *P*_*5*_ = 0.50).

Overall, CO females showed significantly more homotypic (0.61 +/- 0.17) compared to heterotypic (0.39 +/- 0.17) copulations (*P* = <0.0001, Figure 1c), with no heterogeneity between replicate populations (*P* = 0.12, Table S3). CO_*5*_ females showed an overall significant deviation from the null hypothesis, with more homotypic compared to heterotypic copulations (*P* = <0.001). CO_*1*_ and CO_*3*_ populations showed no overall deviation from a random mating ratio of 1:1 (*P*_*1*_ = 0.10, *P*_*5*_ = 0.07). When data was pooled across trials, both CO_*3*_ and CO_*5*_ populations showed significant deviation from a random mating ratio (*P*_*3*_ = 0.02, *P*_*5*_ = <0.0001). There was no significant heterogeneity among trials within any of the populations (*P*_*1*_ = 0.10, *P*_*3*_ = 0.58, *P*_*5*_ = 0.96).

### Post-mating prezygotic reproductive isolation

#### Hatchability

Cross identity had a significant effect on egg hatchability (*P* = <0.01, Table S4). The random effect of replicate population was not significant (*P* = 0.89). The only significant differences in hatch rate was the decline of CO_*i*_♀ × A_*i*_♂ F2 hybrids compared to ACO and CO parental lines (Figure 2a). Overall, hatchability was high with average values between 0.96 +/- 0.02 (CO_*i*_♀ × A_*i*_♂ F2 hybrids) and 0.98 +/- 0.01 (ACO parentals).

**Figure 2.**
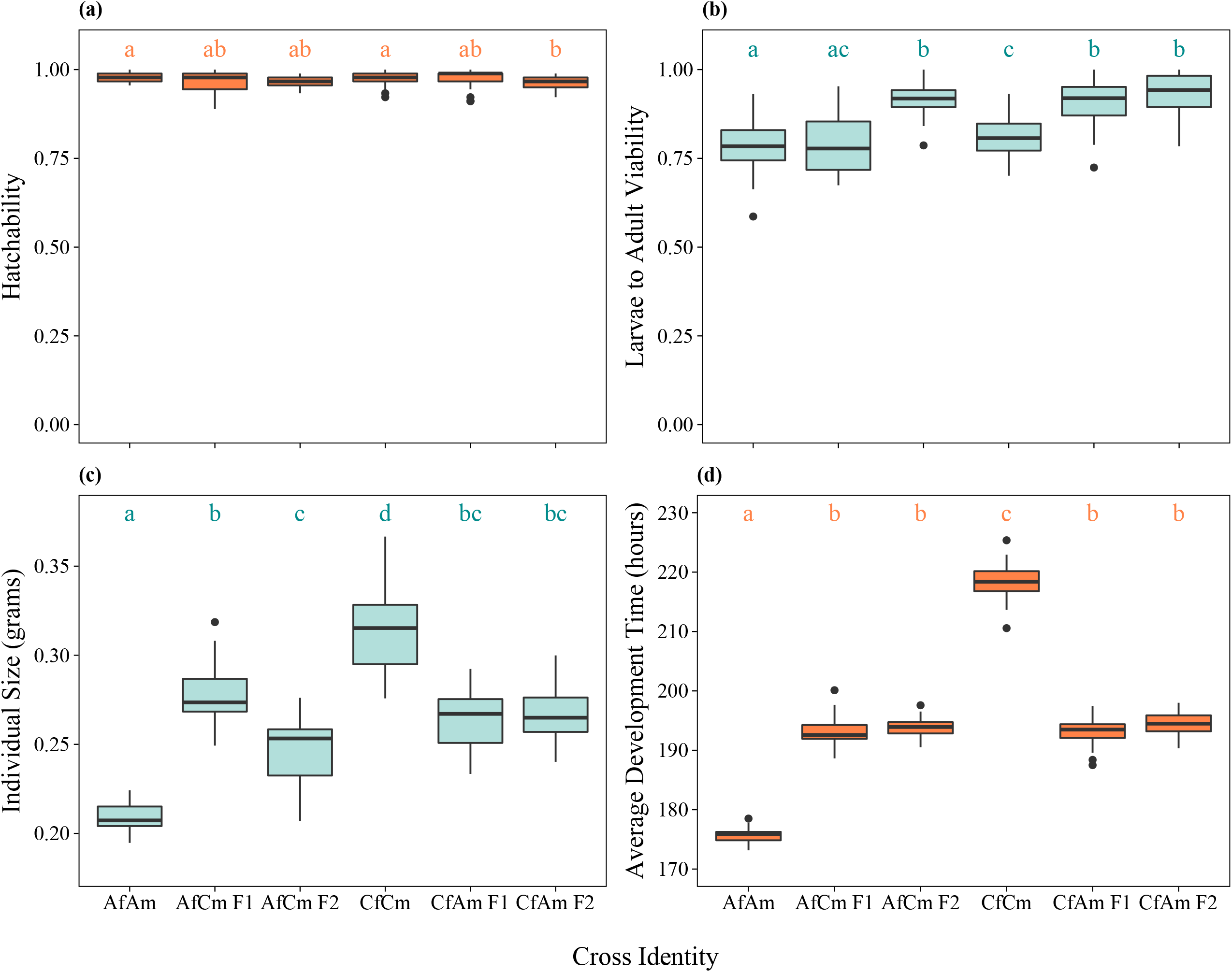
The hatchability (a), egg to adult viability (b), body weights (c) and average development time (d) of flies from parental and hybrid crosses. Letters indicate significant differences between cross identities.

### Postzygotic reproductive isolation

#### Larvae to adult viability

There was also a significant effect of cross identity (*P* = <0.0001, Table S4) and replicate population (*P* = <0.01) on the viability of flies. On average, larvae to adult viability was lowest for ACO parental (0.78 +/- 0.08) and highest for CO_*i*_♀ × ACO_*i*_♂ F2 flies (0.93 +/- 0.06). Both CO_*i*_♀ × ACO_*i*_♂ hybrids and ACO_*i*_♀ × CO_*i*_♂ F2 hybrids demonstrated significantly higher viability than ACO, CO and ACO_*i*_♀ × CO_*i*_♂ F1 flies and CO flies showed significantly higher viability than ACO flies (Figure 2b)..

#### Development time

There was a significant effect of cross identity (*P* = <0.0001, Table S5), but not replicate population (*P* = 0.51) on the development time of flies. All four hybrid crosses demonstrated development times that were significantly longer than ACO parental flies (175.76 +/- 1.33 hours) and significantly shorter than CO parental flies (218.32 +/- 3.22 hours). There were no significant differences in development times among the four hybrid crosses (Figure 2d).

#### Body size

Cross identity also had a significant effect on the body size of flies (*P* = <0.0001, Table S5). The random effect of the replicate population was not significant (*P* =0.23). All four hybrid lines showed average individual weights that were significantly higher than parental ACO lines (0.21 +/- 0.01 g), and significantly lower than parental CO lines (0.31 +/- 0.02 g). Hybrid ACO_*i*_♀ × CO_*i*_♂ F1 flies were also significantly heavier than ACO_*i*_♀ × CO_*i*_♂ F2 flies (Figure 2c).

#### Reproductive fitness

Cross identity had a significant effect on female reproductive fitness (*P* = <0.0001, Table S6). Parental CO lines showed the highest average fitness (0.24 +/- 0.12), parental ACO lines the lowest (0.12 +/- 0.10) and hybrids displayed intermediate fitness between parental lines (Figure 3). All cross identities produced significantly more offspring than parental ACO females and flies from all cross identities, except for CO_*i*_♀ × ACO_*i*_♂ F1 females, produced significantly fewer offspring than parental CO flies. In both cases, second generation (F2) females displayed lower reproductive fitness than first generation (F1) hybrids. There was no significant effect of population replicate on female fertility (*P* = 1).

**Figure 3.**
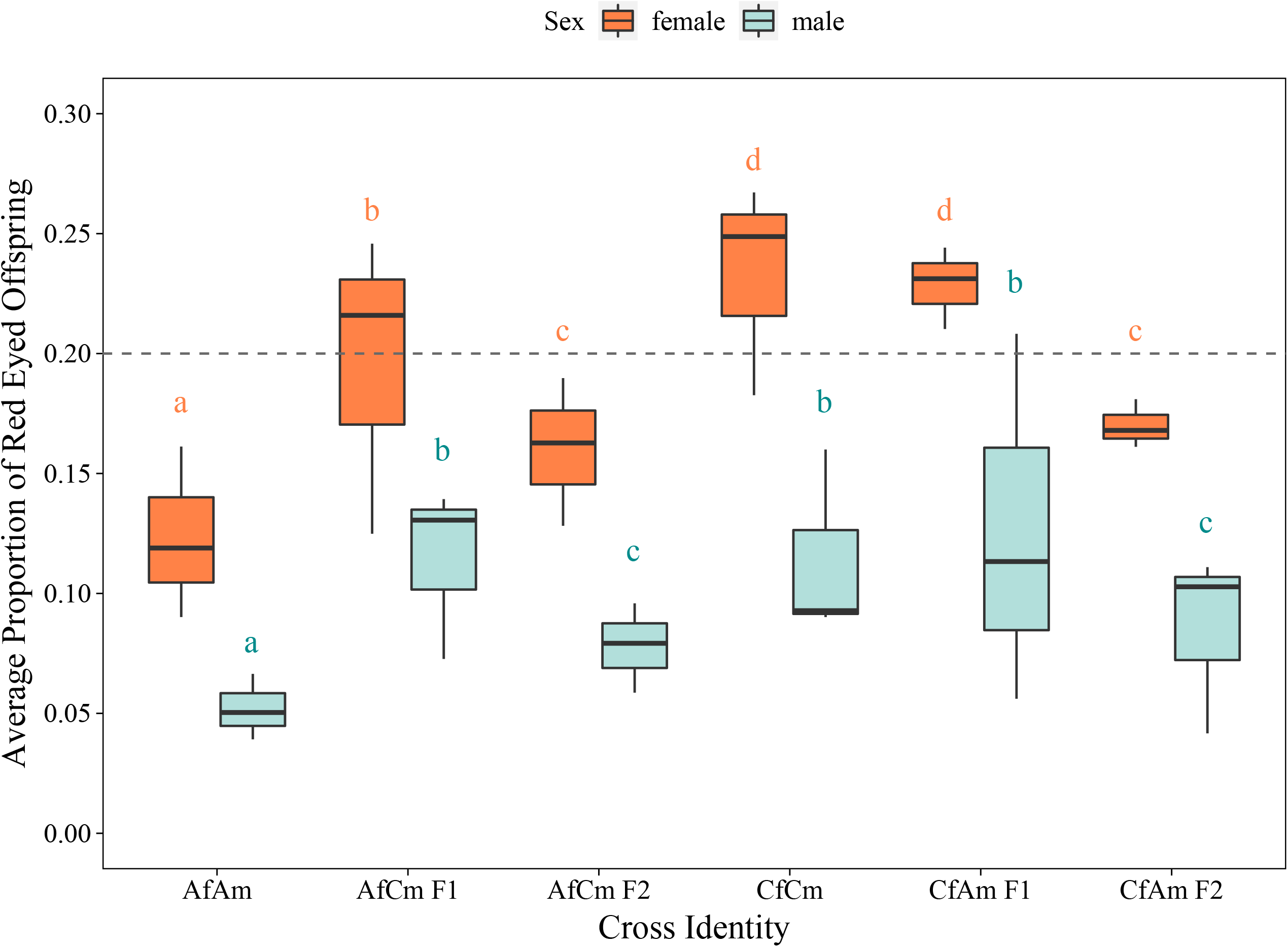
The average proportion of red eyed offspring produced by flies from parental and hybrid crosses, in competition with brown eyed competitors. Average proportions were determined for each replicate population and plotted. The dashed line indicates the expected proportion of ⅕ of the offspring. Letters indicate significant differences between cross identities.

There was a significant fixed effect of cross identity (*P* = <0.0001, Table S6) and random effect of replicate population (*P* = 0.02) on the fertility of male flies. CO_*i*_♀ × ACO_*i*_♂ F1 male hybrids showed the highest average fitness (0.13 +/- 0.20) and ACO males the lowest (0.05 +/- 0.11). F1 hybrid and CO males demonstrated a significantly higher fitness than F2 and ACO males (Figure 3). F2 hybrid males sired significantly more offspring than A males. Similar to that observed in the female assay, male F2 hybrids displayed significantly lower fitness than F1 hybrids. While female flies displayed fitness values around the expected value, males of both parental and hybrid cross identities produced a lower proportion of offspring than expected when in competition with four competitors (Figure 3).

## DISCUSSION

Our experimental assays show the striking differentiation between the ACO and CO populations after over three decades and 1 500 generations of divergent selection on demography. The ACO population adults eclose nearly two days earlier than the CO population adults do. The clear tradeoff between speed and size is evident, as A adults are about 33% lighter than CO adults, and this is likely related to the strong differences in adult fitness we observed between the two selection treatments. Interestingly, the tradeoff between developmental speed and juvenile viability (Chippindale et al. 1997) is not apparent here or in other recent unpublished data on these populations, suggesting compensatory evolution over time. Given these differences in morphology and life history, we asked if we could detect the early stages of the speciation process for these populations in allopatry under different ecological circumstances.

To summarise our results, we found evidence for the evolution of premating barriers to gene flow between these diverged populations. Mate-choice assays generally reflected positive assortative mating patterns driven by female mate-choice. Females from both focal populations showed strong homotypic mate choice in individual tests, but this pattern was less clear in group tests, particularly as the smaller A females tended towards heterotypic mates. We show that postmating prezygotic barriers have not evolved to the point of detectability between these populations in the form of gamete incompatibility, measured through egg hatchability. We also find no evidence of postzygotic reproductive isolation in the form of survival to reproductive maturity, and measurements of development time and body size show that hybrids display intermediate trait values relative to the parental treatments. We observed only suggestive trends for postzygotic reproductive isolation in the form of hybrid breakdown in competitive reproductive fitness assays. F_1_ hybrids display intermediate fitness to that of the two parentals, however, there is a tendency for F_2_ hybrid offspring to display lower fitness than the F_1_ hybrids; while potentially an indicator of evolving Dobzhansky-Muller incompatibilities, our data are merely suggestive at this juncture, meriting further investigation.

It has been suggested that prezygotic isolating barriers will typically evolve before postzygotic barriers, and *Drosophila* seems to adhere to this rule (Coyne & Orr 2004, Arthur & Dyer 2015). Within the category of prezygotic barriers, premating RI is expected to evolve before postmating-prezygotic RI (Turissini et al. 2018). Our data are generally in accord with these predictions, as our diverged population pairs show marked premating reproductive isolation, at least under individual choice tests, but clear evidence for later stage RI is lacking.

Strong divergence in body size exists between ACO and CO animals (Xiao et al. 2019; data herein) and are presumed to correlate with differences in persistence/resistance capabilities between the selection treatments. This size difference led us to expect the presence of some premating reproductive isolation. However, we predicted premating isolation in one direction, on the basis of greater resistance from larger (CO) females, greater persistence of CO males, and a possible preference for large mates. That directional RI was noted by Ghosh & Joshi (2012) in a study similar to ours: bigger mates were preferred by both large/slow and small/fast flies. Contrary to that precedent, females of both regimes displayed strong homotypic mate choice in individual tests. This result is especially intriguing for the ACO females, as they displayed a preference for smaller ACO males even though *D. melanogaster* females are expected to prefer larger males (Pitnick 1991, Jagadeeshan et al. 2015, but see Prasad et al. 2007).

Male mate choice tests may present trickier interpretations. ACO selected males largely mated with homotypic females, but male COs mated nearly randomly. While ACO males appear to be showing positive assortative mating preferences, both types of males were observed making repeated courtship attempts with both homotypic and heterotypic females (unquantified observations) suggesting that the skewed mating pattern could be the result of the increased capability of CO females to resist unwanted mating attempts, particularly from the smaller A males. In populations of flies evolved under a similar pair of life history selection regimes, Ghosh and Joshi (2012) found that small males attempted to court large females, but were unsuccessful, and, as a result, mated more frequently with small females. We therefore cannot easily ascribe mating assortment to the choice of the male, who may be undiscriminating under the test conditions, particularly with no competition for mates. A-type females are also under tremendous pressure to mate, as they are selected to lay eggs about 24h after becoming sexually mature. In the absence of competition, many males may have simply taken the most receptive mate first; for the CO males this may have counterbalanced a homotypic preference, creating the ambivalent outcome observed. The results of the mate choice assays taken together suggest that premating isolation in this system is most strongly driven by female choice.

Body size is a likely trait involved in the evolution of reproductive isolation between the population pairs tested. However, it remains possible that the patterns of assortative mating stem from traits that are yet to be characterised. Some possibilities include courtship behaviours (mating call, wing movements) and cuticular hydrocarbon profiles that may have diverged under the divergent selection regimes. Besides female preference, body size differences also play a clear role in the differential capacities displayed by females to resist mating attempts by males, which creates a further reproductive barrier between large females and small males. Despite large body size differences between the selection treatments, it seems unlikely that the patterns of positive assortative mating can be explained by mechanical incompatibilities in copulation alone; some earlier anecdotal observations suggested they may exist, but we saw no clear evidence in these trials.

In the group mate choice assays, female CO flies continued to show homotypic mating preference, but A females displayed reversed preferences, mating more frequently with heterotypic males than homotypic males. The existence of heterotypic mate choice in groups, in spite of homotypic mate choice in individual assays suggests that the nature of the assay plays a role in the mating patterns observed. In the group mating assays, vials were more crowded (32 individuals vs. 3) - potentially creating circumstances where resisting forcible mating-attempts is more difficult for females. This is a likely explanation given that only the small A females displayed reversed mate choice preferences, while the larger CO females largely reproduced the same mate choice patterns as displayed in individual assays. At the time these experiments were conducted (since changed) there was an important difference in the mating environment of the ACO and CO selection treatments: whilst the early-reproducing ACO’s primarily mated in vials, with only a brief phase in cages for egg-laying, the CO’s, with 14 days in cages probably do all mating relevant to fitness in the cage environment. With these differences in the “natural” mating environment of our LTEE, the mate choice patterns found in individual assays seem to show us that A-selected females have a preference for homotypic mating but are unable to exercise it in a group mating setting with larger males.

Asynchronous development time alone can contribute to reproductive isolation, as seen in the experimental work of Miyatake and Shimuzu (1999) on the melon fly. Because the mating times diverged in lines selected for development time, significant RI arose that was evidenced in experimental mate choice trials. Similarly, the extreme early reproduction regime of the A populations means that neither CO individuals nor A/CO hybrids could hybridize, as the latter groups would not even be sexually mature before a generation is completed. This scenario is, of course, artificial because natural populations generally do not grow on synchronised, discrete generations. they are not directly responsible for the assortative mating patterns observed in the controlled mate-choice assays.

Post-mating, we report no evidence of reproductive isolation deriving from gamete incompatibility, as would be evident in reduced egg hatch rates. Similarly, our measurements of larva to adult viability data show that there are no hybrid treatments with lower levels of viability than the purebred parental populations. In fact, we observed trends of hybrid vigour, with some of the hybrid treatments displaying higher larva to adult viability than the parental treatments. Interestingly, the larva to adult viabilities in F2 treatments were significantly higher than that of the parental selection treatments and one F1 treatment. While the increased viability in hybrids could suggest the existence of homozygous deleterious variation in developmental loci in our selection regimes, this should be consistently seen in both F1 and F2 generations, instead of being more common in the F2 generation. Development time and body size displayed no suggestion of abnormalities indicative of developmental instability, with intermediate hybrid trait values relative to the parental treatments in both sexes.

The competitive reproductive fitness of hybrids was generally intermediate to the parental treatments, and we found no evidence of hybrid sterility. Given that both assays subjected a focal fly to 4 competitors of the same sex, we expect focal flies to sire/dam ⅕th of the offspring produced, if all else was equal. In female reproductive fitness assays, our populations did cluster around the predicted threshold, with hybrids scoring intermediate fitness levels relative to the low fitness ACOs and high fitness COs. In male reproductive fitness assays though, fitness fell well below the predicted threshold in all treatments. We rule out the possibility of this being a case of Haldane’s rule, because male fitness falls well below predicted levels in parental males as well as hybrids. We believe that this is a consequence of the usage of IV*b* flies as competitors/mates in these assays. Selected as common ancestors to both selection regimes, the IV*b*s have evolved in allopatry with both the populations, with a slightly different maintenance regime. We believe that the reduced male fitness estimates may be a consequence of homotypic mate choice by IV*b* females in male fitness assays. While we do see different levels of competitive reproductive output for the different treatments in the female assay, it is not possible to disentangle male mate choice from innate fecundity and harm resistance differences in the females of different selection regimes. Overall, in spite of the depressed male fitness output relative to predictions, hybrids in both sexes still consistently showed intermediacy in fitness, suggesting no reproductive sterility and a link to an additive trait such as body size.

The comparison between F1 and F2 reproductive fitness revealed an interesting trend: F1 fitness was consistently higher than that of F2 offspring, however, this difference was statistically significant in only 2 out of 12 comparisons (2 reciprocal crosses*3 replicate treatment pairs*2 sexes). We interpret this with caution as a possible trend of hybrid breakdown as a consequence of incompatible gene complexes evolving in the two selection regimes.

We generally found high repeatability across our three independent replicates, which suggests that the characters measured and pre-mating RI observed stem from consistent treatment differences between the CO and ACO populations. Internal random processes of divergence in allopatry, such as the coevolution of arbitrary signaller-receiver relationships in allopatry may also drive the evolution of RI. A system like ours opens the door to a potential test for such divergence if RI were investigated between replicate populations *within* selection treatment, rather than *between* treatments. Although our three decade experiment seems too young to see more than hints of post-mating RI, more sensitive queries of characters like developmental stability (e.g., as measured by fluctuating asymmetry) may be of interest. Further investigation of the sources of premating RI and its environmental sensitivity are warranted.

