## Supplementary material for "Evolution of reproductive isolation in a long-term evolution experiment with *Drosophila melanogaster*: 30 years of divergent life history selection": Table

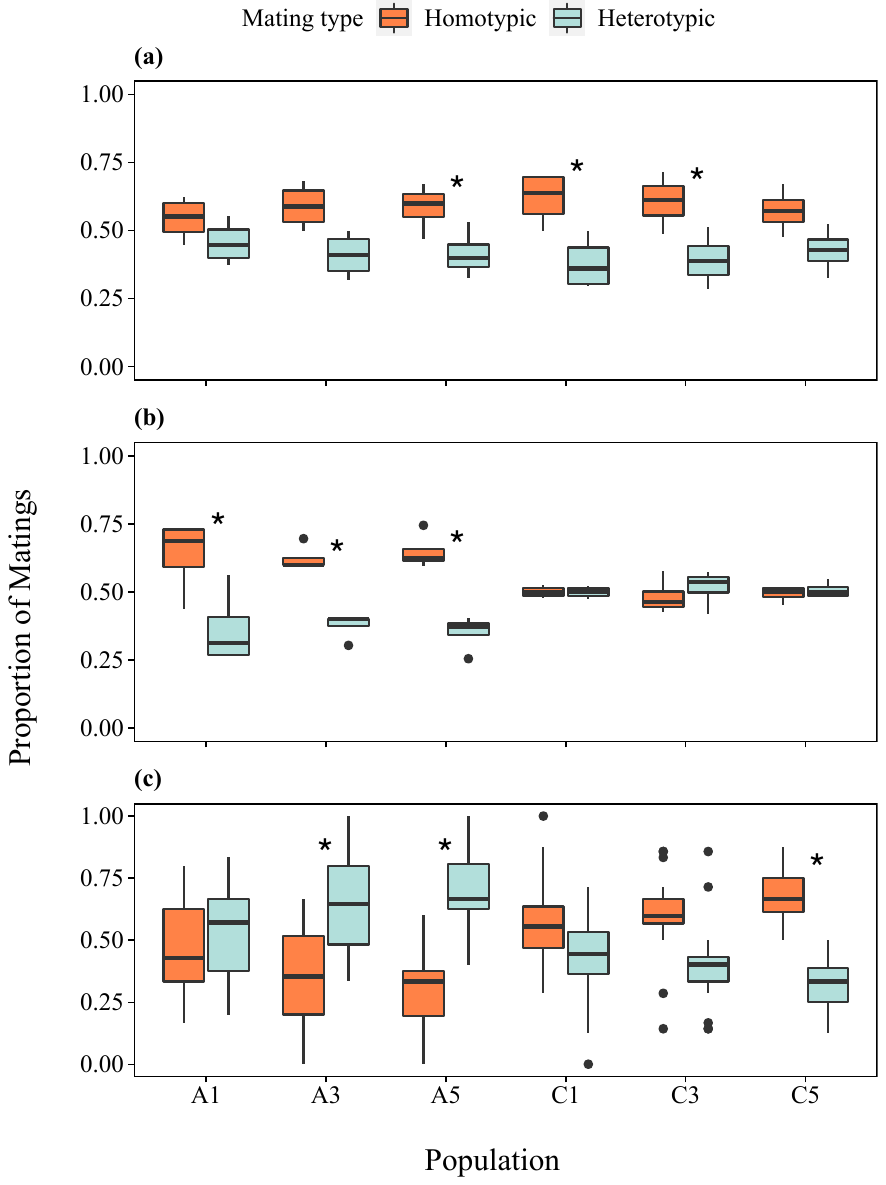
**Figure S1.** The proportion of homotypic compared to heterotypic matings recorded during the female (a), male (b), and group (c) mate choice assay for each replicate population. An asterisk represents a significant deviation from a random mating ratio as indicated by a repeated G-test for goodness of fit (p<0.05).

**Table S1.** The results of the repeated G-test for goodness of fit for the female mate choice assay.

| **Population** | **Heterogeneity G** | **df** | **P-value** | **Pooled G** | **df** | **P-value** | **Total G** | **df** | **P-value** |
| --- | --- | --- | --- | --- | --- | --- | --- | --- | --- |
| ***A1*** | *3.7* | *3* | *0.30* | *1.55* | *1* | *0.21* | *5.25* | *4* | *0.26* |
| ***A3*** | *3.87* | *3* | *0.28* | *5.40* | *1* | *0.02* | *9.27* | *4* | *0.05* |
| ***A5*** | *4.54* | *3* | *0.21* | *5.47* | *1* | *0.02* | *10.02* | *4* | *0.04* |
| ***Overall A*** | *0.82* | *2* | *0.66* | *11.59* | *1* | *<0.001* | *12.42* | *3* | *0.01* |
| ***C1*** | *5.68* | *3* | *0.13* | *11.13* | *1* | *<0.001* | *16.81* | *4* | *< 0.01* |
| ***C3*** | *5.4* | *3* | *0.15* | *9.16* | *1* | *<0.01* | *14.56* | *4* | *0.01* |
| ***C5*** | *3.87* | *3* | *0.28* | *4.42* | *1* | *0.04* | *8.29* | *4* | *0.08* |
| ***Overall C*** | *0.85* | *2* | *0.65* | *23.86* | *1* | *<0.00001* | *24.71* | *3* | *< 0.0001* |

^1^ A repeated G-test was conducted on the level of each replicate population, then on the level of the overall selection regime. A total p-value of less than 0.05 indicates significance.

| **Population** | **Heterogeneity G** | **df** | **P-value** | **Pooled G** | **df** | **P-value** | **Total G** | **df** | **P-value** |
| --- | --- | --- | --- | --- | --- | --- | --- | --- | --- |
| ***A1*** | *11.62* | *3* | *0.01* | *13.61* | *1* | *<0.001* | *25.22* | *4* | *<0.0001* |
| ***A3*** | *1.42* | *3* | *0.70* | *11.37* | *1* | *<0.001* | *12.79* | *4* | *0.01* |
| ***A5*** | *3.11* | *3* | *0.38* | *17.85* | *1* | *<0.0001* | *21.11* | *4* | *<0.001* |
| ***Overall A*** | *0.41* | *2* | *0.81* | *46.85* | *1* | *<0.00001* | *47.27* | *3* | *<0.00001* |
| ***C1*** | *0.23* | *3* | *0.97* | *0* | *1* | *1* | *0.23* | *4* | *0.99* |
| ***C3*** | *2.18* | *3* | *0.54* | *0.29* | *1* | *0.59* | *2.47* | *4* | *0.65* |
| ***C5*** | *0.55* | *3* | *0.91* | *0.05* | *1* | *0.83* | *0.6* | *4* | *0.96* |
| ***Overall C*** | *0.15* | *2* | *0.93* | *0.19* | *1* | *0.67* | *0.34* | *3* | *0.95* |

**Table S2.** The results of the repeated G-test for goodness of fit for the male mate choice assay.

^1^A repeated G-test was conducted on the level of each replicate population, then on the level of the overall selection regime. A total p-value of less than 0.05 indicates significance.

| **Population** | **Heterogeneity G** | | **df** | **P-value** | **Pooled G** | **df** | **P-value** | **Total G** | **df** | **P-value** |
| --- | --- | --- | --- | --- | --- | --- | --- | --- | --- | --- |
| ***A1*** | | *0.48* | *1* | *0.49* | *0.01* | *1* | *0.92* | *0.49* | *2* | *0.78* |
| ***A3*** | | *0.58* | *1* | *0.45* | *7.64* | *1* | *0.01* | *8.23* | *2* | *0.02* |
| ***A5*** | | *0.46* | *1* | *0.50* | *18.75* | *1* | *<0.0001* | *19.21* | *2* | *<0.0001* |
| ***Overall A*** | | *8.87* | *2* | *0.01* | *17.53* | *1* | *<0.0001* | *26.4* | *3* | *<0.00001* |
| ***C1*** | | *2.75* | *1* | *0.10* | *1.8* | *1* | *0.18* | *4.55* | *2* | *0.10* |
| ***C3*** | | *0.3* | *1* | *0.58* | *5.13* | *1* | *0.02* | *5.43* | *2* | *0.07* |
| ***C5*** | | *0.002* | *1* | *0.96* | *16.21* | *1* | *<0.0001* | *16.21* | *2* | *<0.001* |
| ***Overall C*** | | *4.31* | *2* | *0.12* | *18.83* | *1* | *<0.0001* | *23.14* | *3* | *<0.0001* |

**Table S3.** The results of the repeated G-test for goodness of fit for the group mate choice assay.

^1^A repeated G-test was conducted on the level of each replicate population, then on the level of the overall selection regime. A total p-value of less than 0.05 indicates significance.

Table S4. Full GLMM results for the hatchability and larval to adult viability of flies from parental and hybrid crosses.

| *Response* | *Factor Type* | |  | *χ^2^* | *df* | *P* |
| --- | --- | --- | --- | --- | --- | --- |
| Hatchability | Fixed | **Cross Identity** |  | ***19.10*** | ***5, 154*** | ***<0.01*** |
|  |  |  | *AIC* | *χ^2^* | *df* | *P* |
|  | Random | Replicate Population | 645.96 | 0.02 | 1 | 0.89 |
|  |  |  |  | *χ^2^* | *df* | *P* |
| Larvae to Adult Viability | Fixed | **Cross Identity** |  | **431.53** | **5, 154** | **<0.0001** |
|  |  |  | *AIC* | *χ^2^* | *df* | *P* |
|  | Random | **Replicate Population** | **1212.40** | **10.08** | **1** | **<0.01** |

^1^Bold font indicates statistical significance (P < 0.05). GLMMs followed the general format: Response ~ Cross Identity + (1|Replicate Population), family = binomial.

Table S5. Full LMM results for the development time and body size of flies from parental and hybrid crosses.

| *Response* | *Factor Type* | | *MS* | *df* | *F* | *P* |
| --- | --- | --- | --- | --- | --- | --- |
| Development Time | Fixed | **Cross Identity** | ***4974.90*** | ***5, 154*** | ***987.18*** | **<0.0001** |
|  |  |  | *AIC* | *Df* | *LRT* | *P* |
|  | Random | Replicate Population | 731.97 | 1 | 0.43 | 0.51 |
|  |  |  | *MS* | *df* | *F* | *P* |
| Body Size | Fixed | **Cross Identity** | ***0.01*** | **5, 64** | **41.36** | **<0.0001** |
|  |  |  | *AIC* | *Df* | *LRT* | *P* |
|  | Random | Replicate Population | -304.10 | 1 | 1.41 | 0.23 |

^1^Bold font indicates statistical significance (P < 0.05). The LMMs followed the formula:

Response ~ Cross Identity + (1|Replicate Population).

Table S6. Full GLMM results for the fertility of female and male flies from parental and hybrid crosses.

| *Sex* | *Response* | *Factor Type* | |  | *χ^2^* | *df* | *P* |
| --- | --- | --- | --- | --- | --- | --- | --- |
| Female | Proportion Red Eyed | Fixed | **Cross Identity** |  | ***794.57*** | ***5, 515*** | ***<0.0001*** |
|  |  |  |  | *AIC* | *χ^2^* | *df* | *P* |
|  |  | Random | Replicate Population | 9165.5 | 0 | 1 | 1 |
|  |  |  |  |  | *χ^2^* | *df* | *P* |
| Male | Proportion Red Eyed | Fixed | **Cross Identity** |  | **472.05** | **5, 502** | **<0.0001** |
|  |  |  |  | *AIC* | *χ^2^* | *df* | *P* |
|  |  | Random | **Replicate Population** | **13576** | **5.08** | **1** | **0.02** |

^1^Fertility was determined as the proportion of red eyed offspring produced in competition with brown eyed competitors. Bold font indicates statistical significance (P < 0.05). GLMMs followed the general format: Proportion Red Eyed ~ Cross Identity + (1|Replicate Population), family = binomial.
